# Feedback-mediated signal conversion promotes viral fitness

**DOI:** 10.1101/361071

**Authors:** Noam Vardi, Sonali Chaturvedi, Leor S. Weinberger

## Abstract

A fundamental signal-processing problem is how biological systems maintain phenotypic states (i.e., canalization) long after degradation of initial catalyst signals. For example, to efficiently replicate, herpesviruses (e.g., human cytomegalovirus, HCMV) rapidly counteract cell-mediated silencing using trans-activators packaged in the tegument of the infecting virion particle. But, the activity of these tegument trans-activators is inherently transient—they undergo immediate proteolysis but delayed synthesis—and how transient activation sustains lytic viral gene expression despite cell-mediated silencing is unclear. By constructing a two-color, conditional-feedback HCMV mutant, we find that positive feedback in HCMV’s Immediate Early 1 (IE1) protein is of sufficient strength to sustain HCMV lytic expression. Single-cell time-lapse imaging and mathematical modeling show that IE1 positive feedback converts transient transactivation signals from tegument pp71 proteins into sustained lytic expression, which is obligate for efficient viral replication, whereas attenuating feedback decreases fitness by promoting a reversible silenced state. Together, these results identify a regulatory mechanism enabling herpesviruses to sustain expression despite transient activation signals—akin to early electronic transistors—and expose a potential target for therapeutic intervention.

## INTRODUCTION

A common problem in information-transmission systems is how to robustly maintain signal strength despite the inevitable signal decay that occurs through any transmission medium. For example, in the last century, the need to re-amplify telephone signals, which diminished in strength over transmission-line distance, spurred innovations, such as vacuum-tube repeaters and the point-of-contact and junction transistors (1, 2). In developmental biology, a corresponding robustness problem regards how developmental states are stably maintained, often referred to as ‘canalization’ for how the ‘canal’ walls in an epigenetic landscape stabilize the developmental state (3). During canalization, biological systems must overcome the signal-decay problem and stably maintain a developmental state long after degradation of initiating signals and various mechanisms evolved to convert transient signals to sustained outputs (4-9). The fundamental issue facing biological signaling molecules, often proteins or nucleic acids, is that they have intrinsic degradation times mediated either by environmental mechanisms (e.g., ultraviolet radiation) or active intracellular machinery (e.g., the proteasome). The mechanisms allowing cells to sustain differentiated phenotypes long after degradation of the initial (transient) differentiation signals, remain actively studied (5, 7).

Viruses face a version of the canalization problem as they balance a temporal tradeoff: on the one hand, a virus must *rapidly* counteract cellular defenses to initiate its lytic cycle, but on the other hand, the virus must sustain repression of cellular defenses over the course of its intracellular viral lifecycle, which can last for many hours or even days. To antagonize cellular defenses as rapidly as possible after infection, herpesviruses have evolved to carry trans-activator molecules within the infecting virion particle and thereby circumvent the delays inherent to *de novo* gene expression. However, these trans-activator molecules are immediately subject to cellular degradation mechanisms upon infection, and herpesvirus lifecycles can extend for days (10). How, or if, these trans-activator signals are sustained across the course of the viral lifecycle remains unknown. Beta herpesviruses, having intracellular lifecycles lasting over 4 days, appear to have a particularly large transient-versus-sustained signaling problem to overcome.

The beta herpesvirus human cytomegalovirus (HCMV)—a leading cause of birth defects and transplant failures—must rapidly overcome innate host-cell defense mechanisms to initiate its lytic cycle. Cellular promyelocytic leukemia nuclear bodies (PML-NBs; a.k.a. nuclear domain 10, ND10) (11)—containing PML, the death domain-associated protein (DAXX) and SP100—act to repress viral gene expression and prevent HCMV from initiating its lytic cycle (12-14). To counteract these cellular defenses, HCMV virions package the viral transactivator phospho-protein 71 (pp71) in virion’s tegument layer (15). pp71 is the major tegument transactivator protein and interacts with DAXX to promote its degradation. Upon infection, pp71 tegument proteins are released into the cell, disrupt PML-NBs to antagonize the cellular silencing machinery, and activate the HCMV Major Immediate-Early Promoter (MIEP). The MIEP initiates the viral transcription program and its main protein products, the Immediate Early 1 (IE1) and 2 (IE2) proteins are obligate for lytic expression by regulating host and viral genes (16-19), manipulating the host cell cycle (20, 21), and circumventing host innate defense (22, 23); whereas IE2 is obligate at all MOIs, IE1 appears to be obligate only at low MOI (11, 24-26). We asked how HCMV is able to sustain the pp71 trans-activation signal—which is inherently transient due to pp71 proteolysis—to avoid cell-mediated silencing over an extended 96-hour cellular infection cycle.

Building off theoretical analysis showing that auto-regulatory circuitry (i.e., feedback) can promote cellular ‘memory’ to transient differentiation signals during developmental canalization (3, 5), we hypothesized that positive feedback may enable the virus to extend a transient trans-activation signal. Positive feedback, whereby a signaling molecule auto-activates its own expression or drives a downstream auto-activating loop, can occur through a variety of mechanisms, including transcriptional transactivation, enhanced translation, and even reduced degradation (for a review see (5)). Such feedback motifs are widely used in biological systems to execute cell-fate decisions in both unicellular and multicellular organisms (5, 8, 27-32). HCMV encodes a strong immediate-early negative-feedback loop to maintain IE2 levels below a cytotoxic threshold (33), but whether HCMV encodes an immediate-early positive feedback loop that could extend transient tegument trans-activation signals is unclear. One possibility is that IE1, which can weakly transactivate the MIEP (34), could encode positive-feedback circuitry to temporally extend the duration of MIEP transactivation, but, whether this auto-regulation was strong enough to be physiologically relevant, was not clear.

Here, through the use of mathematical modeling, we develop an assay to probe IE1 positive-feedback strength and its physiological relevance to viral replication. To test the feedback-mediated temporal extension hypothesis, we constructed a tunable, two-color HCMV reporter virus where both IE1 and IE2 can be followed in real time and IE1 proteolysis can be pharmacologically tuned using a small molecule via a degron tag (35). Using single-cell time-lapse imaging (Movies S1 and S2), we find that IE1 positive feedback is sufficiently strong to convert transient pp71 transactivation into sustained lytic expression. The results show that IE1 positive feedback is essential for sustained MIEP expression and that attenuating the viral positive-feedback strength causes a severe fitness defect by promoting entry to a silenced state. These findings suggest that transcriptional positive feedback in herpesviruses acts as a transistor-like mechanism to temporally extend intrinsically transient tegument signals and facilitates canalization of the productive (lytic) viral expression state.

## RESULTS

### The HCMV major tegument trans-activator has an 8-hour lifetime but modeling predicts that a putative positive-feedback loop could sustain trans-activation

To understand how HCMV can sustain expression from the MIEP despite stable cellular silencing (Figure 1A), we first measured the degradation rate of the major tegument trans-activator pp71 (24) after infection. ARPE-19 cells were synchronously infected with a recombinant HCMV encoding an EYFP-pp71 fusion (36) and followed for 12 hours using time-lapse fluorescence microscopy. pp71 appeared as distinct fluorescent foci within cells with the number of foci decreasing during the first few hours of infection (Figure 1B). Quantifying the temporal kinetics of the number of pp71 foci per cell results in a measured pp71 half-life of 8.1±1.1 hours (Figure 1B), which agrees with cycloheximide-based estimates of the decay rate of total cellular pp71 (Figure S1A-B). Moreover, we were unable to detect cleavage products of the EYFP-pp71 fusion protein in infected cells, indicating that EYFP signal decay likely reflects pp71 decay (Figure S1C). Nevertheless, the 8-hour pp71 half-life we measured was used as an upper limit, since it is possible that fusion of EYFP to pp71 increases its stability (36).

**Figure 1:**
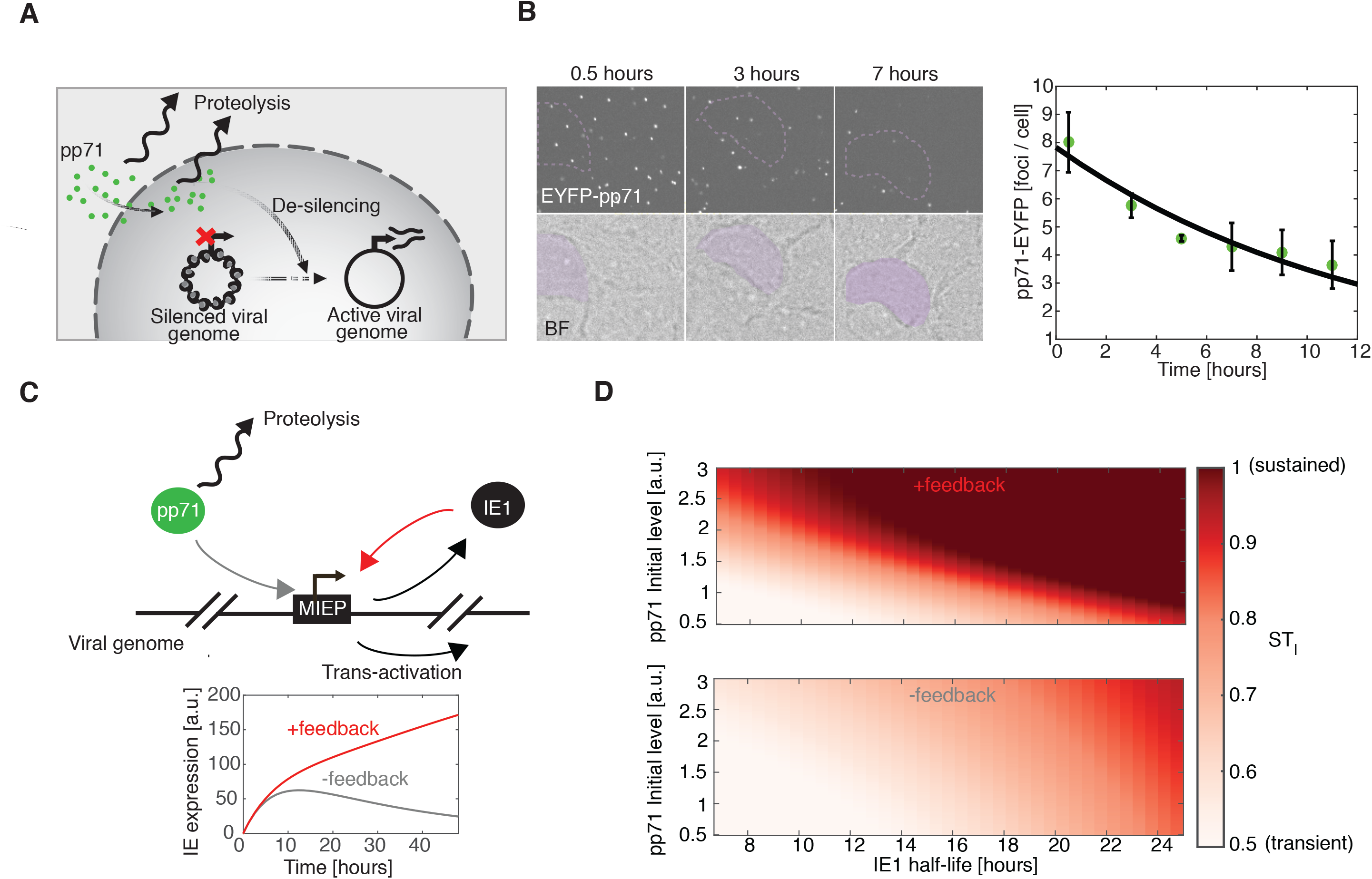
Tegument-derived pp71 is transient in cells, but modeling predicts that positive feedback could sustain transactivation. (*A*) Schematic of initial regulatory events occurring during infection of cells with HCMV. (*B*) Time-lapse fluorescence microscopy of ARPE-19 cells synchronously infected with an EYFP-pp71 HCMV virus (AD169) at MOI of 1 and quantification of pp71 cellular levels over the first 10 hours of infection (an average of 162 cells). (*C-D*) A mathematical model describing pp71 and IE1 dynamics. Equations 1 and 2 were solved numerically, using ODE45 solver (MATLAB^TM^, MathWorks). *ST_I_* was calculated for every set of parameters as the ratio between IE1 expression at 24 hours after infection and maximal IE1 expression. For parameters, see Table S2.

We next used mathematical modeling to simulate the expected MIEP activity given this measured pp71 decay rate. Since IE1 is transcribed from the MIEP during the first few hours of infection, and similar to pp71, IE1 localizes to PML bodies to antagonize these cellular anti-viral defenses (22, 37, 38), we hypothesized that IE1 may play a role in sustaining MIEP expression. To examine the theoretical validity of this hypothesis (Figure 1C), we constructed a highly simplified Ordinary Differential Equation (ODE) model of IE1 expression in the presence of a decaying pp71 trans-activation signal using a ‘course-grained’ modeling approach to generate the following non-linear ODE model:

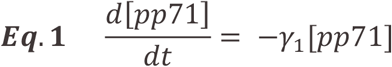

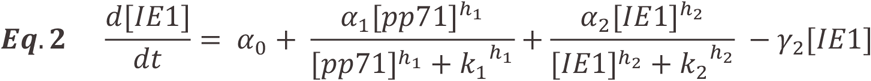

Importantly, the course-grained modeling approach is not intended as a comprehensive description of all known molecular interactions. Instead, the goal is to find a minimum set of interactions and components capable of generating testable predictions about the duration of IE expression—for this reason, the IE2 negative-feedback accelerator circuitry (33) was determined to be expendable (See Figure S2, Table S3). In this highly simplified, course-grained model, only ‘active’ pp71 and IE1 protein levels are considered, and we attempt to limit the number of unmeasured parameters. The pp71 decay, per-capita rate *γ_1_*, is measured above, and the per-capita IE1 proteolysis rate *γ_2_* was previously measured (39). The model also accounts for IE transactivation by pp71 and IE1 (Eq. 2); *α_0_* represents the basal rate of MIEP-driven IE1 expression, *α_1_* represents the transactivation of the MIEP by pp71, *α_2_* accounts for IE1 transactivation of the MIEP and is the putative positive-feedback gain parameter. *k_1_* and *k_2_* are Michaelis-like constants for transactivation, and *h_1_* and *h_2_* are Hill coefficients to account for self-cooperativity of transactivation. Simulations of a more detailed model, which includes the IE2 negative-feedback accelerator circuit (33), yield similar results and qualitatively match the temporal dynamics of IE2 expression observed in single cells (Figure S2).

As expected, simulations of Eqs. 1–2 show that the temporal duration of IE expression is significantly affected by the IE1 degradation rate (Figure 1C). When the IE1 half-life is long (~24 hours in the wild-type case as measured below), IE expression is sustained and monotonically increasing over the first 24 hours. However, when IE1 half-life is reduced, IE expression becomes transient with a dramatic drop during the first 24 hours post-infection. When positive feedback is removed by abrogating IE1 transactivation of the MIEP (*α_2_* set to zero), IE expression does not exhibit sustained expression for either long or short IE1 half life (Figure 1C).

To quantify the duration of IE expression, we also defined a sustained-versus-transient IE-expression index, *ST_I_*—the ratio between IE1 expression level at 24 hours post-infection and the maximal IE1 expression peak in an individual cell (essentially a version of the 2^nd^ derivative test, used here as a canalization measure). When a cell’s *ST_I_* is high (>0.5), the cell is able to sustain at least 50% of peak IE-expression levels, whereas when a cell’s *ST_I_* is low (<0.5) the cell is unable to sustain IE expression and it drops below 50% of its peak value. We examined *ST_I_* over a wide range of parameters and as expected, the *ST_I_* was significantly affected by both the IE1 degradation rate and the initial level of pp71 at the time of infection (Figure 1D). These simulations showed that when IE1 degradation is slow, IE expression is sustained (high *ST_I_*) even when initial pp71 levels are low (Figure 1D, upper panel). However, in the absence of IE1 positive feedback (*α_2_* set to zero), IE expression is transient, and a much longer IE1 half-life (i.e., super physiological) is required to sustain IE expression (Figure 1D, lower panel).

These results suggested that IE1 positive feedback acts to extend the duration of pp71 transactivation and proposed an assay for the presence of IE1 positive feedback, based on movement along the horizontal axis in Figure 1D. If IE1 positive feedback is sufficiently strong, reducing the IE1 half-life (right to left in Figure 1D) should switch IE expression profiles (both IE1 and IE2) from sustained to transient (high to low *ST_I_*). In contrast, if positive feedback is weak or absent, reducing the IE1 half-life (within the physiological range) will have little qualitative effect on the sustained-versus-transient expression profiles. In addition, the simulations predicted that high initial levels of pp71 could also sustain IE expression, even in the absence of feedback (Figure 1D).

### Tuning IE1 degradation reveals a positive-feedback loop required to sustain HCMV immediate early gene expression

To test the model predictions on the role of IE1 positive feedback in sustaining IE expression, we generated an IE1-<u>C</u>onditional <u>M</u>utant <u>D</u>ual-<u>R</u>eporter virus (HCMV-IE1-CMDR). First, on the background of a previously characterized TB40e-IE2-EYFP parent virus (33), we fused an mCherry reporter to the N-terminus of IE1 (automatically also tagging IE2) to enable real-time tracking of the immediate early promoter dynamics during infection (Figure 2A–B); we could not detect cleavage products of the mCherry-IE fusion in infected cells (Figure S3A) indicating that mCherry signal likely reflects IE expression levels. Second, to tune the IE1 degradation rate, we fused an FKBP degron tag (35) to the N-terminus of mCherry-IE1. In the presence of the small molecule Shield-1, the IE1 degradation rate is unperturbed by the FKBP degron, but in the absence of Shield-1, the tagged protein is rapidly degraded. This N-terminus tagging approach for IE1 was required since attempts to fuse proteins to the C-terminus of IE1 generated virus with MOI-dependent replication defects (unpublished data). In addition to the N-terminus IE1 tags, this new reporter virus has EYFP fused to the C-terminus of IE2, so that the IE2 protein can be tracked in parallel to IE1. Since IE1 is far more abundant than IE2 (39) and has a substantially longer half-life, mCherry primarily reports on IE1, with minimal contribution from IE2. In the presence of Shield-1, this IE1-CMDR virus grows with similar replication kinetics to the parent virus (Figure S3B).

**Figure 2:**
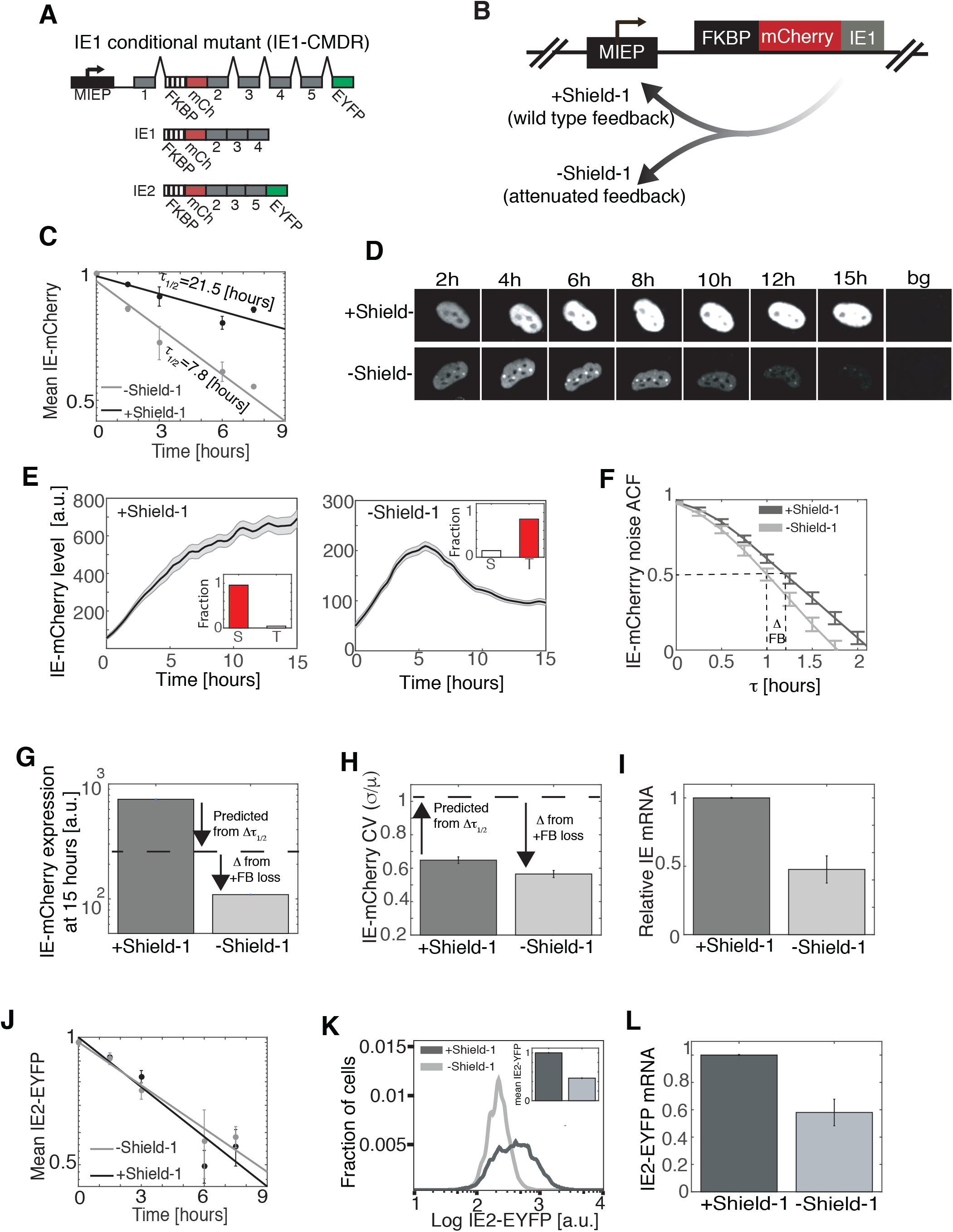
IE1 positive feedback sustains expression from the HCMV immediate early promoter. (*A*) Map of the MIE region of the recombinant HCMV TB40/E IE1 conditional-mutant dual-reporter virus (IE1-CMDR) showing the MIEP (black), IE region exons 1–5 (grey), along with the recombinant fusions: exon 2 is genetically fused at its N terminus to mCherry (mCh; red) and the FKBP degron tag (hashed) that destabilizes the IE1 protein in the absence of the small molecule Shield-1; exon 5 also contains a fusion to EYFP (green) at its C terminus. Translation typically begins at the 5’ end of exon 2 (now FKBP). See also Figure S3. (*B*) Schematic of the putative effects of Shield-1 on IE1-mediated positive feedback. (*C*) Degradation rate of IE1 in cells infected with IE1-CMDR virus as measured by flow cytometry. Protein synthesis was blocked with cyclohexamide at 24 hours after infection (time=0) and the degradation rate was measured ±1 µM Shield-1. Shown is the average of two repeats, and error bars denote standard deviation. Decay rate was calculated by fitting the data to an exponential decay model (solid lines). (D) Representative images from fluorescence time-lapse microscopy of IE-mCherry expression in cells infected with IE1-CMDR virus. Cells were cultured in medium containing 1 µM Shield-1 (top row) or without Shield-1 (bottom row). Cells were cultured in medium with 1 µM Shield-1 (upper row, wild-type feedback) or without Shield-1 (bottom row, attenuated feedback) and tracked over time. (*E*) IE-mCherry expression of cells infected with IE1-CMDR virus. Cells were cultured in medium with 1 µM Shield-1 (left panel, wild-type feedback, averaged over 119 cells) or without Shield-1 (right panel, attenuated feedback, averaged over 153 cells). Bold line denotes mean (i.e., general trend) of the population with grey shading showing standard error. Cell trajectories were digitally synchronized to the first detection of mCherry signal. Inset: the fraction of cells with sustained (S, *ST_I_* > 0.5) or transient (T, *ST_I_* < 0.5) IE-mCherry expression over three biological repeats. (*F*) High Frequency Autocorrelation function (ACF) of cells infected with IE1-CMDR virus in the presence of 1 µM Shield-1 (dark grey) or without Shield-1 (light grey). Shown is an average over 100 cells each; error bars denote standard error. (*G-H*) IE-mCherry mean expression level and noise (CV) 15 hours after infection. Dashed line is the expected value from change in IE1 half-life alone (Δ from τ_1/2_), whereas the additional difference (Δ) is ascribed to the loss of IE1 positive feedback. (*I*) Relative IE-mCherry mRNA levels (from RT-qPCR) in cells infected with IE1-CMDR virus ±1 µM Shield-1. (*J*) Degradation rate of IE2-EYFP in cells infected with IE1-CMDR virus as measured by flow cytometry. Protein synthesis was blocked with cyclohexamide at 24 hours after infection (time=0) and degradation rate was measured ±1 µM Shield-1. Shown is the average of two repeats, and error bars denote standard deviation. Decay rate was calculated by fitting the data to an exponential decay model (solid lines). (*K*) Flow cytometry for IE2-EYFP levels in cells infected with IE1-CMDR virus. Cells were cultured in medium with 1 µM Shield-1 or without Shield-1. Inset: Normalized mean fluorescence of IE2-EYFP. (*L*) Relative IE2-EYFP mRNA levels (from RT-qPCR) in cells infected with IE1-CMDR virus ±1 µM Shield-1.

To measure the degradation rate of IE1 protein, the decay in mCherry intensity was quantified after cycloheximide translation block in the presence and absence of 1µM Shield-1 at 24 hours after IE1-CMDR infection of ARPE-19 cells. The results show that the FKBP degron decreases IE1 protein half-life from 21.5 to 7.8 hours, or ~three-fold (Figures 2C and S4).

Next, we used single-cell resolution time-lapse microscopy to track IE-expression dynamics in ARPE-19 cells infected with the HCMV IE1-CMDR virus. In cells infected in the presence of 1 µM Shield-1 (+Shield-1), IE gene expression from the MIEP reached steady state ~10 hours after the infection and remained at steady state (*ST_I_* > 0.5) in >95% of cells (Figures 2D,-E and Movie S1). These dynamics are in temporal agreement with previous results using the HCMV AD169 lab strain (33). Strikingly, infection in the absence of Shield-1 (-Shield-1) results in transient IE expression (*ST_I_* < 0.5) of IE1 (Figure 2D–E and Movie S2) in most cells (>80%). This steep drop in IE1 levels in the absence of Shield-1 occurs despite IE expression initiating at a similar time compared to the +Shield-1 condition.

To quantify changes in feedback strength, we used high-frequency autocorrelation function (HF-ACF) analysis. ACF analysis is a common signal-processing technique for time-lapse data that analyzes the frequency of fluctuations, with changes in half-correlation time (ct_50_) of fluctuation used to determine biophysical properties (e.g., in fluorescence spectroscopy) (40, 41). Shifts in the ct_50_ of the HF-ACF are a sensitive reporter of feedback strength and the HFACF is largely unaffected by changes in protein half-life (40, 41). When HF-ACF of the IE1-mCherry single-cell trajectories is analyzed (Figure 2F), it shows that IE1 ct_50_ is significantly reduced when Shield-1 is removed, indicating attenuation of positive feedback (40, 41).

This IE1 positive feedback, in fact, appears to be required to account for the observed changes in IE mean-expression level and variance. Specifically, the observed change in IE1 level when Shield-1 is removed is substantially greater than can be accounted by a three-fold change in IE1 half life (Figure 2G). In the absence of positive feedback, a three-fold decrease in IE1-mcherry half-life should confer a three-fold decrease in IE1-mcherry steady-state levels. However, IE-mCherry levels 15 hours after infection show a seven-fold decrease. This > 3-fold change in expression level is consistent with attenuation of an IE1 positive-feedback loop that was acting to amplify MIEP expression. Likewise, the change in the magnitude of IE expression fluctuations measured by the coefficient of variation (CV) of the trajectories cannot be accounted for by the change in IE1 half-life alone, based on Poisson scaling (Figure 2H); specifically, for a simple birth-death Poisson process, a seven-fold decrease in mean-expression level should correspond to a ~2.65-fold *increase* in CV (42). However, the data show that this seven-fold decrease in mean IE1 expression levels generates a *decrease* in the IE1 CV (Figure 2G). This decrease in IE1 CV despite decreased mean is consistent with attenuation of positive feedback (6).

To directly test if IE1 destabilization influences MIEP expression, we quantified IE mRNA (as opposed to protein) in the presence and absence of Shield-1 by RT-qPCR (Figure 2I). We observed a ~2-fold decrease in MIEP activity upon Shield-1 removal (Figure 2I), indicating that reduction in IE1’s half-life confers a change in IE mRNA levels, hence an attenuation of the positive feedback strength. This two-fold change in IE mRNA levels accounts for the missing quantitative difference (“Δ”) in the IE protein levels unaccounted for by the IE1 half-life change (Figure 2G).

We also analyzed the affect of IE1 destabilization on IE2 levels. Notably, despite the FKBP tag also being present on the IE2 protein, the degradation rate of IE2 protein does not appear affected by Shield-1 removal (Figure 2J), presumably due to IE2’s relatively short native half-life (33). Despite IE2-EYFP half-life not being affected by the FKBP degron tag, IE2-EYFP expression exhibits a 2-fold decrease in abundance upon infection in the absence of Shield-1 compared to infection in presence of Shield-1 (Figures 2K and S5D). This 2-fold change in IE2 protein levels corresponds to an ~2-fold decrease in IE2-EYFP mRNA levels (Figure 2L). We also analyzed the HF-ACF for IE2-EYFP—whose half-life remains unchanged by Shield-1—and found a ct_50_ shift similar to the shift in IE1 ct_50_ (Figure S5), further validating the presence of an IE1 positive-feedback loop.

Overall, these single-cell imaging data, of both IE1 and IE2, indicate the presence of an IE1 positive-feedback loop that is attenuated when IE1 half-life is reduced by Shield-1 removal.

### Attenuating IE1 positive feedback decreases viral fitness by promoting entry to a reversible silenced state

Next, to determine if sustained IE expression is required for efficient virus replication and, correspondingly, if transient IE expression results in a replicative fitness defect, we measured HCMV viral replication in the setting of both wild-type IE1 feedback (+Shield-1) and attenuated IE1 feedback (-Shield-1). ARPE-19 cells were infected with the IE1-CMDR virus (MOI=0.02) in the presence of Shield-1 (wild-type feedback) and absence of Shield-1 (attenuated feedback), and resulting virus titers assayed over time by TCID-50. The viral replication kinetics revealed a ~10-fold reduction in viral titer when IE1 positive feedback was attenuated (-Shield-1), compared to the wild-type IE1 feedback case (+Shield-1) (Figure 3A). Although these data do not suggest the presence of positive feedback (which is shown in Figure 2), they indicate that sustained IE expression confers replicative fitness for HCMV.

**Figure 3:**
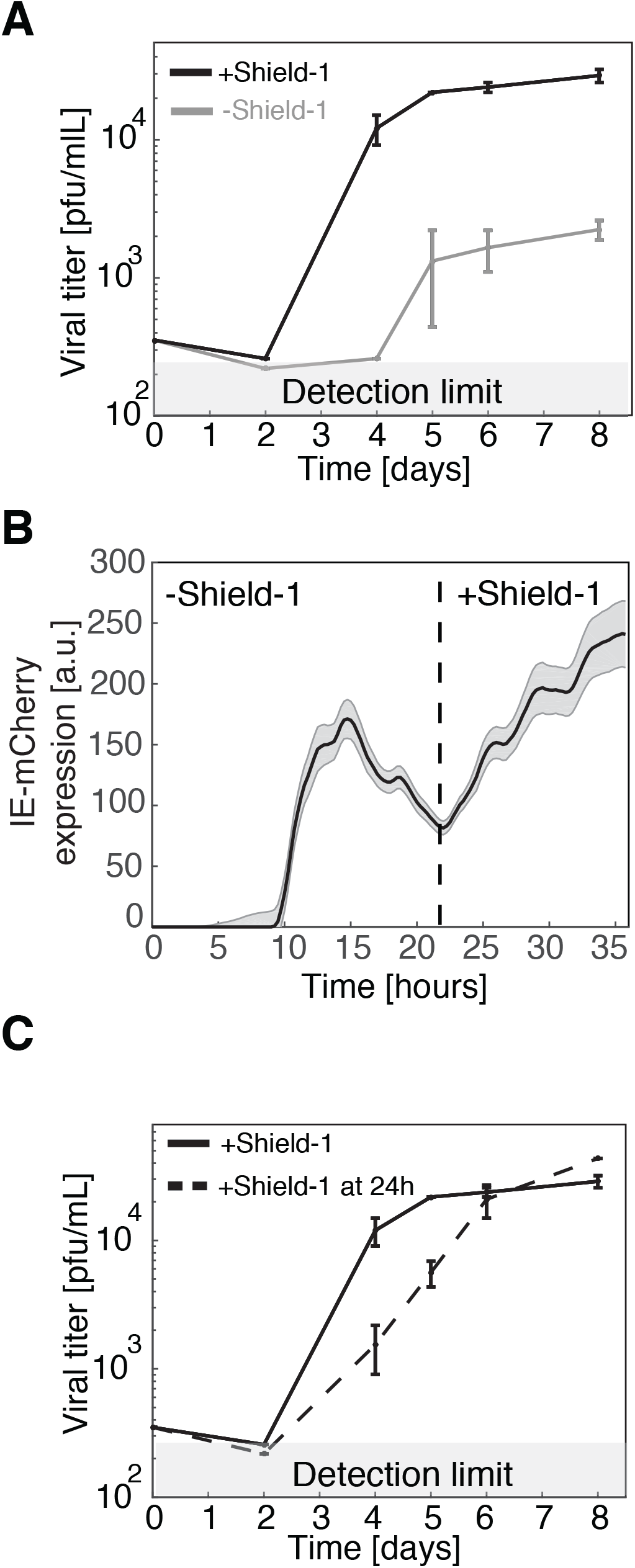
Attenuated IE1 positive feedback decreases replication fitness, but feedback can be rescued to restore fitness. (*A*) Viral titers from cells infected with IE1-CMDR virus. Cells were cultured in media with 1 µM Shield-1 (black, wild-type feedback) or without Shield-1 (grey, attenuated feedback). Virus was harvested at different time points after infection. Viral titers were measured in medium containing 1 µM Shield-1, error bars denote standard deviation. (*B*) IE-mCherry expression from time-laps single-cell microscopy of cells infected with IE1-CMDR virus. Cells were cultured in medium without Shield-1 for 24 hours, and then supplemented with 1 µM Shield-1 for 12 hours. Bold line denotes mean mCherry expression (6×7 cells); grey shading denotes standard error. (*C*) Viral titers from cells infected with IE1-CMDR virus. Cells were cultured in medium with 1 µM Shield-1 (solid line), or Shield-1 was added to the medium after 24 hours (dashed line). Virus was harvested at different time points after infection, titers were measured in media containing 1 µM Shield-1, and error bar denotes standard deviation.

We next asked if the IE-expression transient led to abortive infection or instead promoted a latent-like, silenced state that could be reactivated if IE expression was rescued at a later time. If Shield-1 was added back 24 hours after infection, IE expression kinetics could be restored to a sustained trajectory (Figure 3B), and correspondingly, viral replication could be rescued to wild-type levels (Figure 3C).

### High pp71 abundance partially rescues attenuation of the IE1 positive-feedback loop

As a further test of the model, we examined the second prediction of Eqs. 1–2 that high initial abundance of pp71 could compensate for weak IE1 positive feedback to enable sustained IE expression. To test this prediction, we artificially increased the pp71 tegument load in HCMV virion particles by packaging the IE1-CMDR virus on a pp71-expressing cell line (24). We reasoned that ectopic expression of pp71 from the cell line in addition to the pp71 expressed from the viral genome would result in packaging of more pp71 into virions. Comparing pp71 levels between the two purified viral preparations (pp71^hi^ and pp71^wt^) by western blot showed about a three-fold increase in pp71 abundance in the pp71^hi^ virus isolate (Figure 4A).

**Figure 4:**
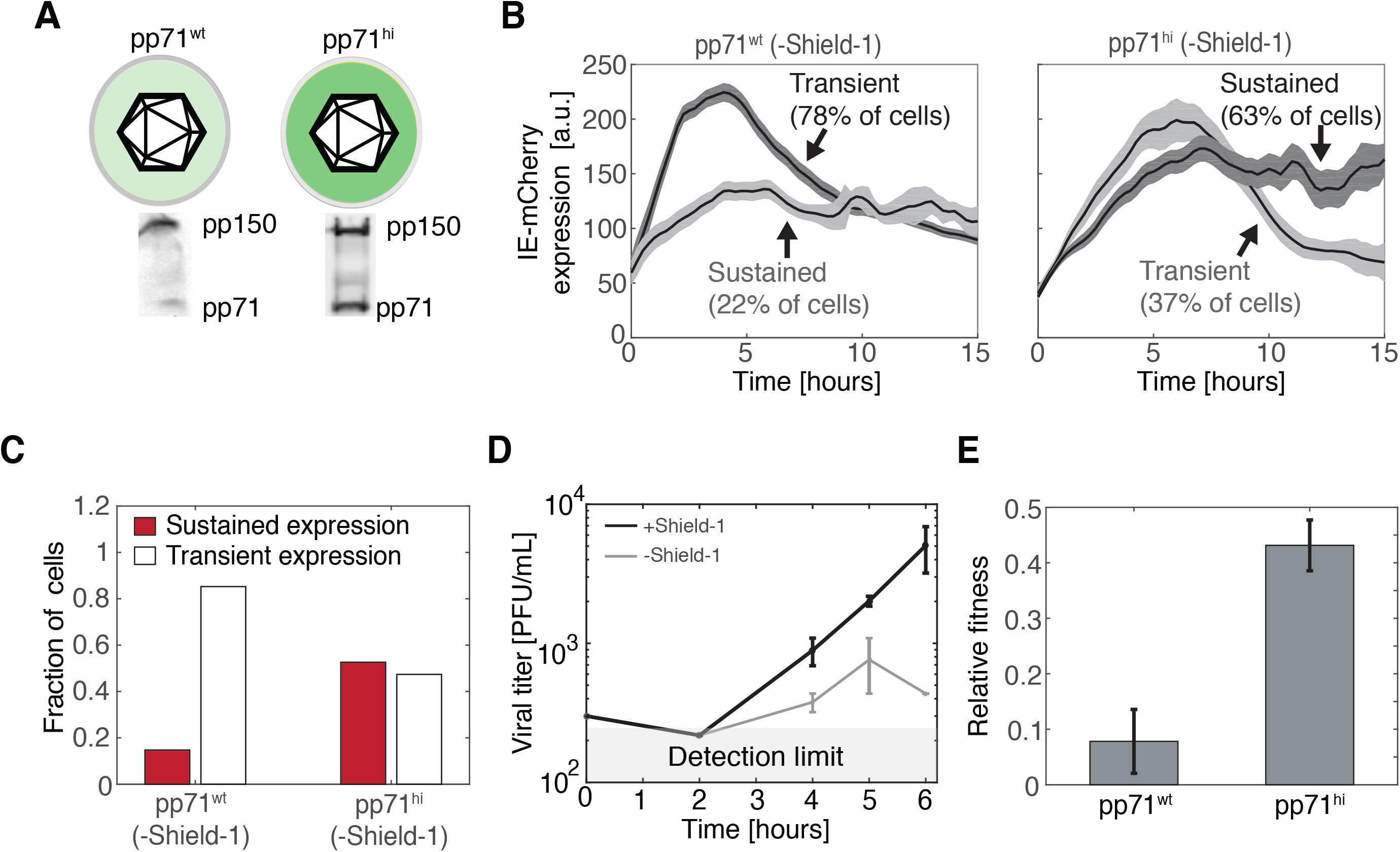
Increased pp71 abundance partially rescues expression and fitness of attenuated feedback virus. (*A*) Western blot analysis of pp71 in viral particles packaged on cells overexpressing pp71 (IE1-CMDR-pp71^hi^) or on non-overexpressing cells (IE1-CMDR-pp71^wt^), (*B*) Representative time-lapse microscopy traces of IE-mCherry expression in cells infected with IE1-CMDR-pp71^hi^ or IE1-CMDR-pp71^wt^. Cells were cultured without Shield-1 (attenuated feedback) and classified as sustained or transient based on expression kinetics as defined above. Bold line denotes Mean mCherry signal of sustained (dark grey) or transient (light grey) cells; shaded area denotes standard error. Cell trajectories were digitally synchronized to the first detection of mCherry signal. (*C*) The fraction of cells with sustained IE-mCherry expression (*ST_I_* > 0.5) after infection with IE1-CMDR-pp71^hi^ virus (116 cells) or IE1-CMDR-pp71^wt^ virus (243 cells). Cells were cultured without Shield-1 (attenuated feedback). P < 0.001 was calculated using a Fisher Exact test. (*D*) High pp71 can partially compensate the fitness lost when feedback is attenuated. Viral titers from cells infected with IE1-CMDR-pp71^hi^ virus ±Sheild-1. Cells were infected in medium with 1 µM Shield-1 (black, wild-type feedback) or without Shield-1 (grey, attenuated feedback), virus was harvested at indicated times after infection, and titers were measured in medium supplemented with 1 µM Shield-1. Compare to Figure 3A. (*E*) High pp71 partially compensates for fitness lost when feedback is attenuated. Relative single-round fitness (titer at day 4) for either IE1-CMDR-pp71^wt^ (left) and IE1-CMDR-pp71^hi^ (right) when feedback is attenuated (-Shield-1) normalized to viral titer for corresponding wild-type feedback (+Shield-1) case (p=0.041, two-tailed t-test).

Next, we followed IE expression in cells infected with either IE1-CMDR-pp71^hi^ or IE1-CMDR-pp71^wt^ in an attenuated feedback setting (i.e., -Shield-1) using time-lapse microscopy. Strikingly, when cells are infected with the IE1-CMDR-pp71^hi^ virus, the percentage of infected cells exhibiting sustained IE gene expression is >50% (Figure 4B–C), despite the absence of Shield-1 (i.e., attenuated feedback), whereas only ~20% show sustained IE expression when infected with the IE1-CMDR-pp71^wt^ virus in the absence of Shield-1. Thus, the results appear to validate the model prediction that increased pp71 can compensate for attenuated feedback to sustain IE expression kinetics.

Next, we used these results to estimate the relative contribution of IE1 and pp71 to the activation of the MIEP. To this end, we created a stochastic version of the course-grain model (see Table S4, Figure S6 and supplementary information). We varied IE1 and pp71 trans-activation levels and examined which regime of parameters yields an increase in the percentage of stable cells from 20% to 50% when the initial pp71 levels increase by 3-fold. The results suggest that in order to satisfy these constrains, MIEP activation by pp71 should be similar to its activation by IE1 (Figure S6); if auto-activation by IE1 were substantially stronger than pp71 per capita activation, the simulations predict that the probabilities of stable cells would be much greater than experimentally observed.

Finally, we checked if pp71’s partial rescue of sustained IE expression in the attenuated-feedback setting also translated into a rescue of viral fitness. Cells were infected at low MOI (0.02) with the IE1-CMDR-pp71^hi^ virus in the presence or absence of 1 µM Shield-1, (i.e., wild-type feedback and attenuated feedback, respectively), and resulting virus titers assayed over time by TCID-50 (Figure 4D). For the pp71^hi^ infection, viral titers decrease two-fold four days post-infection, when feedback is attenuated (Figure 4E). Compared to the >10-fold decrease in fitness when feedback is attenuated in pp71^wt^ infection, this two-fold decrease in fitness in the pp71^hi^ setting is substantially less (Figure 4E), indicating that boosting initial pp71 levels can partially compensate for attenuated IE1 positive feedback to mitigate the viral fitness defect.

Overall, these results suggest that the duration of IE expression is important for viral replication and that IE1 positive feedback converts inherently transient pp71 transactivation signals into sustained IE expression profiles to drive efficient viral replication.

## DISCUSSION

Herpesviruses face a temporal tradeoff between the need to rapidly counteract cellular defenses and initiate lytic infection and the need to sustain the resulting lytic expression patterns during infection. This balancing appears to have resulted in a transient-signaling dilemma as the most rapid mechanism to initiate transactivation is to circumvent *de novo* gene expression and package mature transactivator molecules directly within the infecting virion particle; but doing so subjects transactivator molecules to degradation (without de novo production) immediately upon infection of the cell. Beta herpesviruses, such as HCMV, must overcome a particularly severe transient-signaling problem since their infection cycle lasts for days, but the half-life of its major transactivation signal (pp71) is hours (Figure 1). Mathematical modeling predicted that a putative positive-feedback circuit encoded by HCMV’s IE1 could sustain HCMV MIEP expression despite pp71 degradation (Figure 1) and suggested an assay to probe for IE1 positive feedback. This assay relied on altering the IE1 degradation rate, which required construction of a conditional-mutant dual-reporter IE1 virus (IE1-CMDR). Time-lapse imaging of IE1-CMDR indicated the presence of IE1 positive-feedback circuitry (Figure 2 and Movies S1–S2) and showed that this IE1 positive-feedback circuit confers replicative fitness on the virus (Figure 3).

One technical caveat is that the pp71 half-life we measured (8.1±1.1 hours) is based on the decay of the EYFP-pp71 fluorescence signal after infection of cells with EYFP-pp71 virus. Since it is possible that fusion of EYFP to pp71 increases pp71 half-life (36), 8 hours serves as an upper limit for pp71 half-life. However, if the pp71 half-life is in fact shorter than 8 hours, the need for a mechanism (e.g. positive feedback) that compensates for pp71 decay is even more pronounced.

Interestingly, the mechanisms of action by which pp71 and IE1 antagonize silencing from the PML bodies are different: pp71 induces SUMOyilation and degradation of DAXX (26), while IE1 blocks PML SUMOylation, leading to dispersion of the PML bodies (43). Nonetheless, the data herein reveal a functional overlap between these different mechanisms of action: while IE1 positive feedback sustains expression from the MIEP during pp71 degradation, boosting pp71 levels in the virion can partially rescue MIEP expression and viral replicative fitness despite attenuated IE1 positive feedback (Figure 4). These data suggest that IE1 activity should reach a certain threshold at early times of infection—in order to facilitate efficient viral replication—and that IE1 might be dispensable at late times of infection, as it can be partially complemented by high levels of pp71 at early times.

A limitation of the model we develop (Eqs. 1–2) is that it focuses exclusively on the early times after infection. At later times, different IE2 isoforms such as IE2 40 and IE2 60 are expressed. While these isoforms are not essential for viral replication they appear to have a contribution to viral fitness as their deletion affects the expression of other viral genes such as pp65 and the DNA replication factor UL84 (19, 44). Hence, it is possible that some of the difference in viral fitness we detect in the absence of Shield-1 (Figure 3A) is due to a decrease in the abundance of these other IE2 isoforms.

Intriguingly, this feedback circuitry bears some resemblance to the latency control circuit in HIV-1, where a decision between proviral latency and active replication is regulated by the Tat transcriptional positive-feedback loop (45). However, there are important mechanistic differences as there is no reported tegument equivalent in HIV-1. Instead the viral integration site of the HIV-1 provirus determines the basal activity of the viral promoter, and Tat directly auto-activates its own promoter (the LTR) without acting through PML disruption. It is also not clear if HIV-1 faces a transient-vs.-sustained signaling tradeoff as in the herpesviruses since the chromatin-based silencing mechanisms in T cells occur on the order of days to weeks, whereas the HIV-1 lifecycle is completed within about 40 hours (46, 47). Nevertheless, the fact that a positive-feedback loop controls the decision between active and latent replication in such an evolutionarily distant virus points to the potential generality of such feedback loops in regulating viral latency establishment.

From an evolutionary standpoint, it is interesting to speculate as to why pp71 didn’t evolve to directly auto-regulate its own expression (i.e., without the need for the additional IE1 circuit ‘node’). A possible answer is that pp71 expression appears to be cytotoxic as ectopic pp71 expression requires a late promoter (24, 48). So, one possibility is that early pp71 *de-novo* expression could result in cytoxicity for the cell and, hence, IE1 evolved as a complementary mechanism to sustain the initiation of lytic gene expression and circumvent such cytotoxicity.

Overall, this study illustrates how positive-feedback circuitry is essential to sustain a viral lytic expression program and the associated fitness of a herpesvirus. These findings suggest a potentially broad mechanism for sustaining gene expression triggered by a transient signal that might extend to other herpesviruses, such as HSV-1 and EBV, that use tegument proteins to overcome host immune defenses and initiate the viral lytic cycle (24, 49, 50). As such, this mechanism may provide a new set of drug targets for development of herpesviruses antivirals.

## MATERIALS AND METHODS

### Cloning of recombinant virus

The recombinant IE1 Conditional Mutant Dual Reporter virus (IE1-CMDR) was constructed from the parent virus TB40/E IE2-EYFP (33) in two steps using a BAC recombineering method utilizing galK selection (51). The galactokinase (galK) gene, driven by em7 prokaryotic promoter, was amplified from a pgalk plasmid (51), using pcr primers 1 and 2 (See Table S1), with homology arms flanking the 5′ end of exon 2 of IE1. The DNA fragment was transformed into SW105 bacterial cells carrying the TB40/e IE2-EYFP bacmid (33). Cells were grown on agar plates with galactose as the only carbon source, and single colonies were picked sequenced. Next, an mCherry coding gene was PCR amplified from a plasmid (Clontech, 632522), using primers 3 and 4 with homology regions flanking the galK gene. The DNA fragment was transformed into SW105 bacterial cells carrying TB40/e IE2-EYFP with the galK cassette upstream of exon 2. Cells were grown on 2-deoxy-glactose plates for negative selection. Single colonies were picked and sequenced to verify that the mCherry was in frame with IE1. To fuse an FKBP degron tag upstream of the mCherry reporter the same process was repeated, this time using primers 5 and 6 for inserting the galK gene upstream of the mCherry reporter and primers 7 & 8 to amplify the FKBP tag from the Ld2GITF plasmid (47) and replace the galK with the degron tag (See primers in Table S1). The resulted IE1-CMDR bacmid was then nucleofected into ARPE-19 cells expanded in medium supplemented with 1 µM Shield-1 (Clontech) to produce the IE1-CMDR virus. The fusion of mCherry and IE1 was validated by Western coimmunoprecipitation (Figure S3C; see below for methods).

The EYFP-pp71 recombinant virus (36) was generously contributed by Thomas Stamminger. The fusion of EYFP and pp71 was validated with Co-immunoprecipitation (Figure S1A).

### Cell culture, viruses and media

ARPE-19 cells (ATCC^®^ CRL-2302™) were grown in DMEM/F-12 50/50 (Corning 10-090-CV) medium supplemented with 10% fetal bovine serum (FBS) and 50 U/ml penicillin-streptomycin at 37°C and 5% CO_2_ in a humidified incubator.

WF28 cells (24) and life-extended human foreskin fibroblasts (HFFs) were grown in DMEM (Corning 10-013-CV) medium supplemented with 10% FBS and 50 U/ml penicillin-streptomycin at 37°C and 5% CO_2_ in a humidified incubator.

Tb40/E-IE1 CMDR virus was expanded on ARPE-19 or WF28 cells in the presence of 1 µM Shield-1 for 3 weeks until 100% CPE was detected. Virus was harvested and then concentrated on a 20% sorbitol cushion (WS28 ultracentrifuge, 20,000 RPM for 2 hours). The pellet was re-suspended in 1 ml of fresh medium and tittered on ARPE-19 cells by TCID-50 to determine concentration of the viral stock.

EYFP-pp71 virus was expanded on HFFs for 3 weeks or until 100% CPE was detected, harvested as described above and tittered on HFFs by TCID-50 to determine concentration of the viral stock.

### Time-lapse microscopy

5*10^4^ ARPE19 cells per well were seeded in an 8-well imaging chamber (Lab-TEK, 155409) and grown to confluency for 3 days. HCMV infections were synchronized on ice (MOI=1), virus was removed after 30 minuets, and plate was placed under the microscope for subsequent imaging. All imaging was performed on an Axiovert inverted fluorescence microscope (Carl Zeiss, Oberkochen, Germany), equipped with a Yokogawa spinning disk, a CoolSNAP HQ2 14-bit camera (PhotoMetrics, Tucson, AZ), and laser lines for 488 nm (40% laser power, 400 ms excitation) and 561 nm (40% laser power, 200 ms excitation). To facilitate time-lapse imaging, the microscope has a programmable stage with definite focus and a stage enclosure that maintains samples at 37°C and 5% CO_2_ with humidity. Images were captured every 15 min. For each position a 5-by-5 X-Y grid was sampled with 5 z-positons at 2.5 µm intervals. The objective used was 40x oil, 1.3NA.

### Cell segmentation and image analysis

For analyzing the MIEP dynamics after infection, a maximal intensity projection was applied to all collected z planes of the same field of view to increase signal intensity. Single-cell tracking and segmentation were performed with custom-written code in MATLAB^TM^ (MathWorks) as described (40). mCherry and EYFP intensities were calculated as the median fluorescence intensity of each segmented cell. Single trajectories were smoothed using a qubic spline (MATLAB) and synchronized in-silico to the first detection of fluorescence to account for variation in MIEP expression start time between cells. Cells that initiated expression more than 12 hours after virus was removed or that tracked for less than 10 hours were discarded from the analysis.

For quantification of pp71-EYFP degradation rate, the number of pp71-EYFP foci were counted at each time point using Imagej^TM^ software by the following steps. A maximal z-projection was applied to all z-stacks planes of the same field of view to increase signal strength, then a uniform threshold was set for all images, and the number of EYFP dots was counted. Cells were stained with DAPI to quantify the number of cells for each field of view.

### HCMV virion purification and western blot

The virus pellet was resuspended in a total volume of 500 µl of phosphate buffered saline (PBS, pH 7.4), and centrifuged at 10,000 rpm for 10 minutes at 4°C to remove cell debris. Supernatant was gently overlaid on a 15–50% sucrose gradient, and centrifuged in SW-41 rotor at 28,000 rpm for 2 hours at 4°C. Three light scattering bands were observed after centrifugation: top, middle and bottom bands contained noninfectious enveloped particles, virions, and dense bodies, respectively (52). The middle band was collected and concentrated by centrifuging for 2 hours at 28,000 rpm with 20% sucrose cushion in SW-41 rotor. The pellet was resuspended in 20 µl PBS buffer and subjected to western blot analysis. Briefly, 5 µl of purified virion sample was added to 1x loading buffer (100 mM tris-HCl, pH 6.8, 200 mM DTT, 4% SDS, 0.1% Bromophenol blue, 20% glycerol), and boiled for 10 minutes at 95°C. Samples and corresponding molecular weight marker (Precision Plus Protein Kaleidoscope Prestained Protein Marker, BIO-RAD) were loaded on 10% SDS-polyacrylamide gel, and ran at 90V for 2 hours in Tris-glycine running buffer (25 mM Tris, 250 mM glycine and 0.1% SDS). The gel was blotted on a PVDF membrane using semi-dry transfer unit at 25V for 30 minutes, and was blocked with 10 ml of Li-cor^TM^ Odyssey blocking buffer for 1 hour at room temperature with gentle shaking. Membrane was treated with primary antibodies, anti-pp71 (1:100 dilution, mouse MAb 2H10-9, gift from Dr. Tom Shenk) and anti-pp150 (1:200 dilution, mouse MAb 36-10, gift from Dr. Bill Britt) for 2 hours at room temperature with gentle shaking. The membrane was than washed three times with wash buffer (1x PBS+0.1% Tween-20), treated with secondary Li-cor^TM^ detection antibody (1:20,000 dilution, goat anti-mouse 800CW), and incubated in dark for 1 hour at room temperature. Membrane was washed three times with wash buffer (1xPBS +0.1% Tween-20) and was scanned on Li-cor Odyssey^TM^ system for detection of pp71 and pp150 protein.

### Co-immunoprecipitation

Infected cells were lysed and subjected to immunoprecipitation using antibodies against mCherry (for IE1-mCherry) andEYFP (pp71-EYFP), where mock is immunoprecipitation of uninfected cells and negative control is immunoprecipitation of virus infected cell extract in the absence of EYFP or mCherry antibody. Briefly, 10^6 cells were lysed in RIPA buffer (50 mM Tris, pH 7.4/150 mM NaCl/1 mM EDTA/1% Nonidet P-40/0.1% SDS/0.5) containing protease inhibitor mixture (Roche), and immunoprecipitated using a mouse antibody against EYFP (anti-GFP antibody, Santa Cruz Biotechnology) to pulldown pp71 and mCherry (anti-mCherry antibody, Abcam) to pulldown IE1. Immunoprecipitations were performed at 4°C for 2 hours. Samples were subsequently pulled down using an anti-mouse IgG (whole molecule)-agarose beads (Sigma-Aldrich) for 1 hour, agarose beads were collected, washed three times in RIPA buffer. Beads were suspended in 10ul of RIPA buffer and was added to 1x loading buffer (100mM tris-HCl (pH6.8), 200mM DTT, 4% SDS, 0.1% Bromophenol blue, 20% glycerol), and boiled for 10 minutes at 95°C. Samples in duplicate and corresponding molecular weight marker (precision plus protein kaleidoscope prestained protein marker) were loaded on 10% SDS-Polyacrylamide gel, and run at 90V for 2 hours in Tris-glycine running buffer (25mM Tris, 250mM glycine and 0.1% SDS). Transfer was carried out by blotting the gel on a PVDF membrane using semi-dry transfer unit (Trans-Blot semi-dry electrophoretic transfer cell, Bio-Rad) at 25V for 30 minutes, and was blocked with 10ml of Li-cor Odyssey™ blocking buffer for 1 hour at room temperature with gentle shaking. Detection as carried out by staining membranes with either anti-pp71 (1:100 dilution, mouse MAb 2H10-9, gift from Tom Shenk) and anti-EYFP primary antibodies (1:1000 dilution, anti-GFP antibody, Santa Cruz Biotechnology) for samples immunoprecipitated against EYFP, or anti-IE1 (1:100 dilution, MAb 810, Chemicon) and anti-mCherry (1:1000 dilution, anti-mCherry antibody, Abcam) for samples immunoprecipitated against mCherry for 2 hours at room temperature with gentle shaking. Membrane were subsequently washed three times with wash buffer (1x PBS+0.1% Tween-20), treated with secondary Li-cor™ detection antibody (1:20,000 dilution, goat anti-mouse 800CW), and incubated in dark for 1 hour at room temperature. Membranes were then washed three times with wash buffer (1x PBS +0.1% Tween-20) and scanned on a Li-cor Odyssey™ system for detection of pp71, mCherry, IE1 andEYFP protein.

### qPCR

ARPE-19 cells cultured were infected with IE1-CMDR virus at MOI = 0.1, cultured in medium with 1 µM Shield-1 or without Shield-1 and harvested 48 hours post infection. Total RNA was extracted using RNEASY RNA isolation kit (QIAGEN, 74104) and reverse transcribed using QuantiTet Reverse Transcription Kit (QIAGEN, 205311). CDNA samples were diluted 1:40 and analyzed on a 7900HT Fast Real-Time PCR System (Thermo-Fisher, 4329003) using designed primers (Table S1) and Fast SYBR Green Master Mix (Applied Biosystems, 4385612). Relative mRNA levels of mCherry and EYFP were quantified using peptidylprolyl isomerase A (PP1A) as a reference gene.

### Flow cytometry and calculation of degradation rates

To quantify IE1 and IE2 degradation rates ARPE-19 cells were seeded in a six-well plate, grown overnight and then infected on ice with IE1-CMDR virus (MOI=0.1) in the presence of 1 µM Shield-1. After infection cells were cultured in fresh medium supplemented with 1 µM Shield-1 for 24 hours. After 24 hours, cells were washed twice with fresh medium and cultured in medium containing 50 µg/ml cyclohexamide (Sigma-Aldrich), to stop protein translation, either with 1 µM Shield-1 or without Shield-1. Cells were harvested at indicated time points and mCherry and EYFP signals measured on an LSRII flow cytometer. Infected cells were defined based on mCherry gating and mean fluorescence intensities of mCherry and EYFP were used to calculate degradation rates by fitting the data to an exponential decay curve (Figure S4).

### Calculations of autocorrelation function

High-frequency noise autocorrelation functions (HF-ACF) were calculated as described in (40) to estimate the change in positive-feedback strength. Positive feedback is predicted to increase the correlation time of the noise HF-ACF by an amount related directly to the feedback strength (40). Derivation of ACFs was done as followed: first, for each cell *m*, fluorescence trajectories were de-trended (normalized) by subtracting from each trajectory the population time dependent average fluorescence (i.e., the general trend), in order to account for gene expression changes that affect all the cells (i.e., high frequency filtering to isolate intrinsic noise). Second, for a given cell *m* the HF-ACF was calculated from:

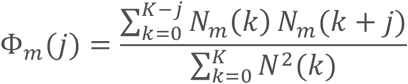

Where *N* denotes to the normalized fluorescence trajectory, and *j* has integer values from 0 to K-1. Next, for *M* cell trajectories the HF-ACF is calculated as:

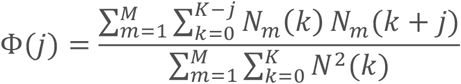

For a full derivation, we refer the reader to (40).

### Mathematical modeling and simulations

The differential equations describing the dynamics of IE1 and pp71 in the main text were simulated using with MATLAB^TM^ (MathWorks) using the ODE45 solver. ST_I_ (Sustained-to-Transient index) was defined as the ratio between IE1 expression at 24 hours post-infection divided by the maximal IE1 expression and examined over a range of parameters. See Tables S2-S3 for parameters values.

The stochastic model of IE1 and pp71 described in Figure S6 and Table S4 was simulated in MATLAB^TM^ by implementing a stochastic Gillespie algorithm (53). For each parameter set, 500 trajectories were simulated over a virtual time-course of 15 hours.

## ACKNOWLEDGMENTS

We thank Thomas Stamminger for the generous contribution of the EYFP-pp71 reporter virus, Cynthia Bolovan-Fritts for support and assistance, Victoria Saykally for technical assistance, and members of our group for fruitful discussions. L.S.W. acknowledges support from the Bowes Distinguished Professorship, the Alfred P. Sloan Research Fellowship, and the NIH Director’s New Innovator Award (OD006677) and Pioneer Award (OD17181) programs.

## SUPPORTING INFORMATION

### SUPPLEMENTARY FIGURES

**Figure S1:**
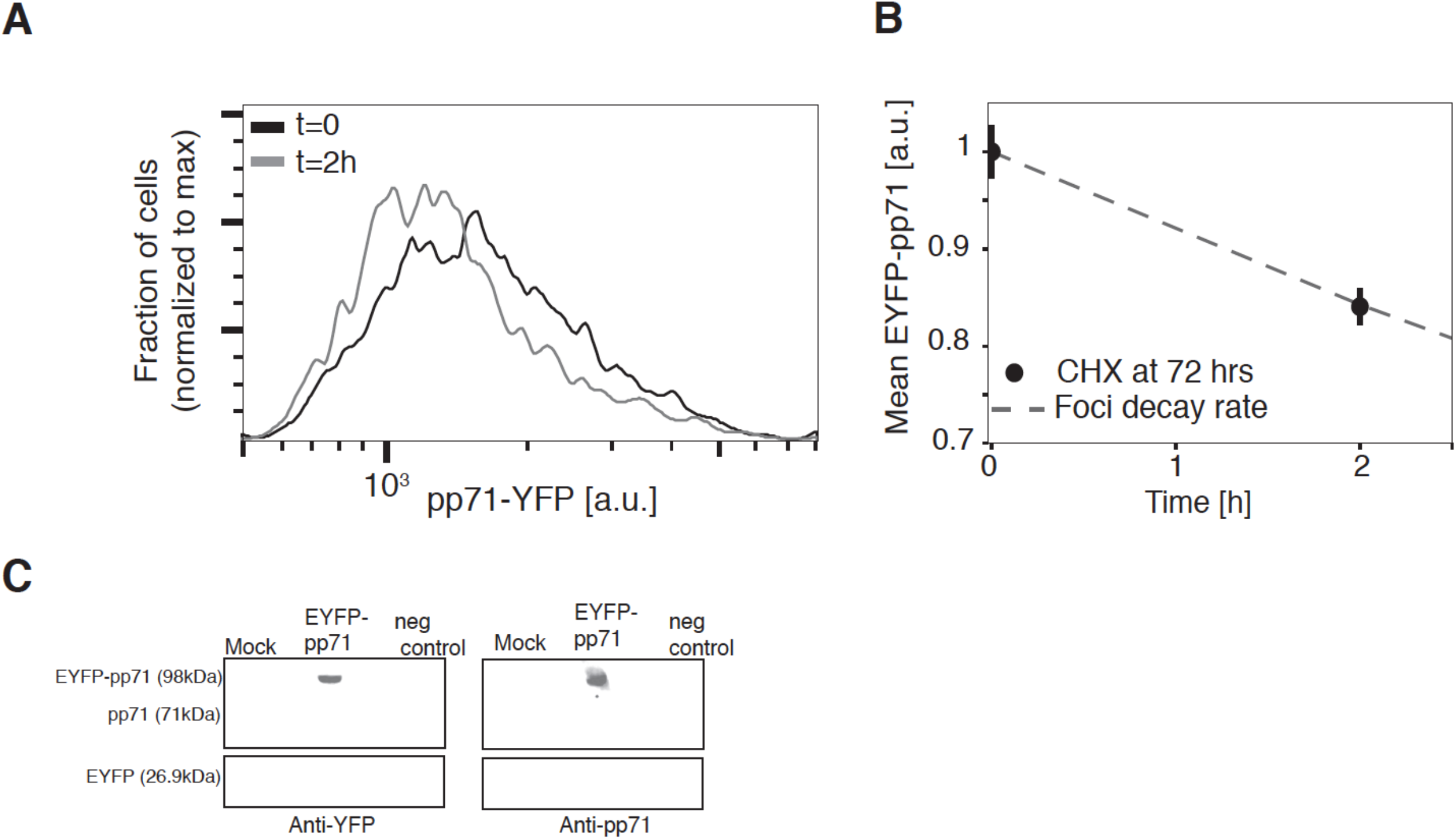
the half-life of EYFP-pp71 is predicted by EYFP-pp71 foci decay rate. (A-B) EYFP-pp71 decay in life-extended human foreskin fibroblasts (HFFs), 72 hours post-infection at MOI = 1, measured by flow cytometry. Data Points are mean fluorescence intensity (average of two repeats, error bars denote standard deviation). Dashed line denotes the decay rate as predicted by EYFP-pp71 foci decay in Figure 1B. (C) Co-ip of pp71 and EYFP shows no free EYFP in infected cells. Cells were either mock infected or infected with EYFP-pp71 virus at MOI = 1, immunoperccipitated with EYFP anti-body and stained with anti-EYFP (left panel) or anti-pp71 (right panel) antibodies. Immunoprecipitation of virus infected cell extract in the absence of EYFP antibody served as a negative control (right columns). See materials and methods for further details.

**Figure S2:**
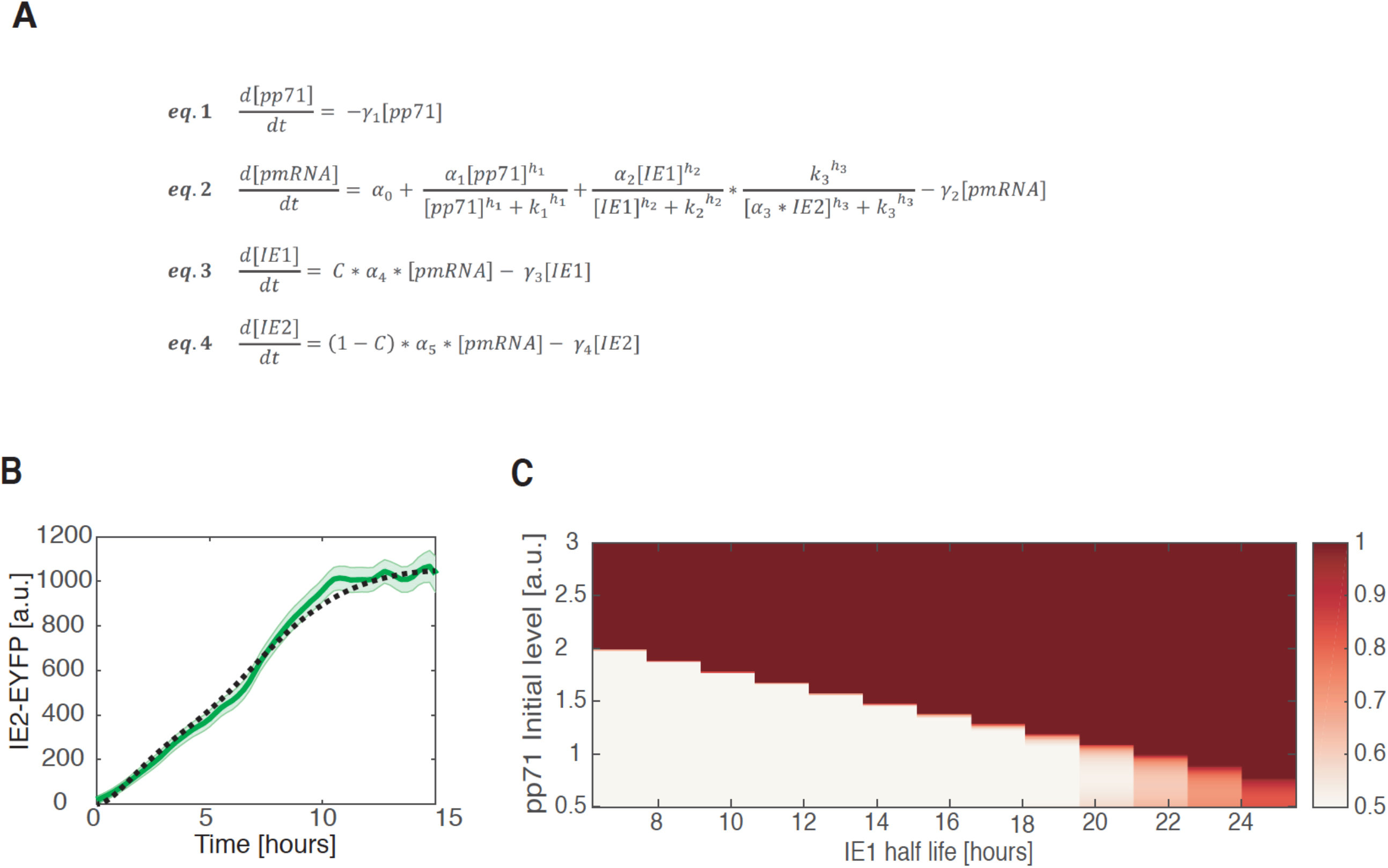
A mathematical model captures the dynamics of IE expression in infected ARPE-19 cells. *(A)* A mathematical model for the dynamics of IE1, IE2 and pp71 after HCMV infection. *(B)* IE2-EYFP expression of cells infected with IE1-CMDR virus. Bold green line denotes mean EYFP expression; shading denotes standard error, n = 119 cells. Numeric simulation of IE2 expression shown as the grey dotted line. *(C)* Sustained-versus-transient IE-expression index (see main text for definition) of the ODE model in panel A as pp71 initial levels and IE1 degradation rate are varied.

**Figure S3:**
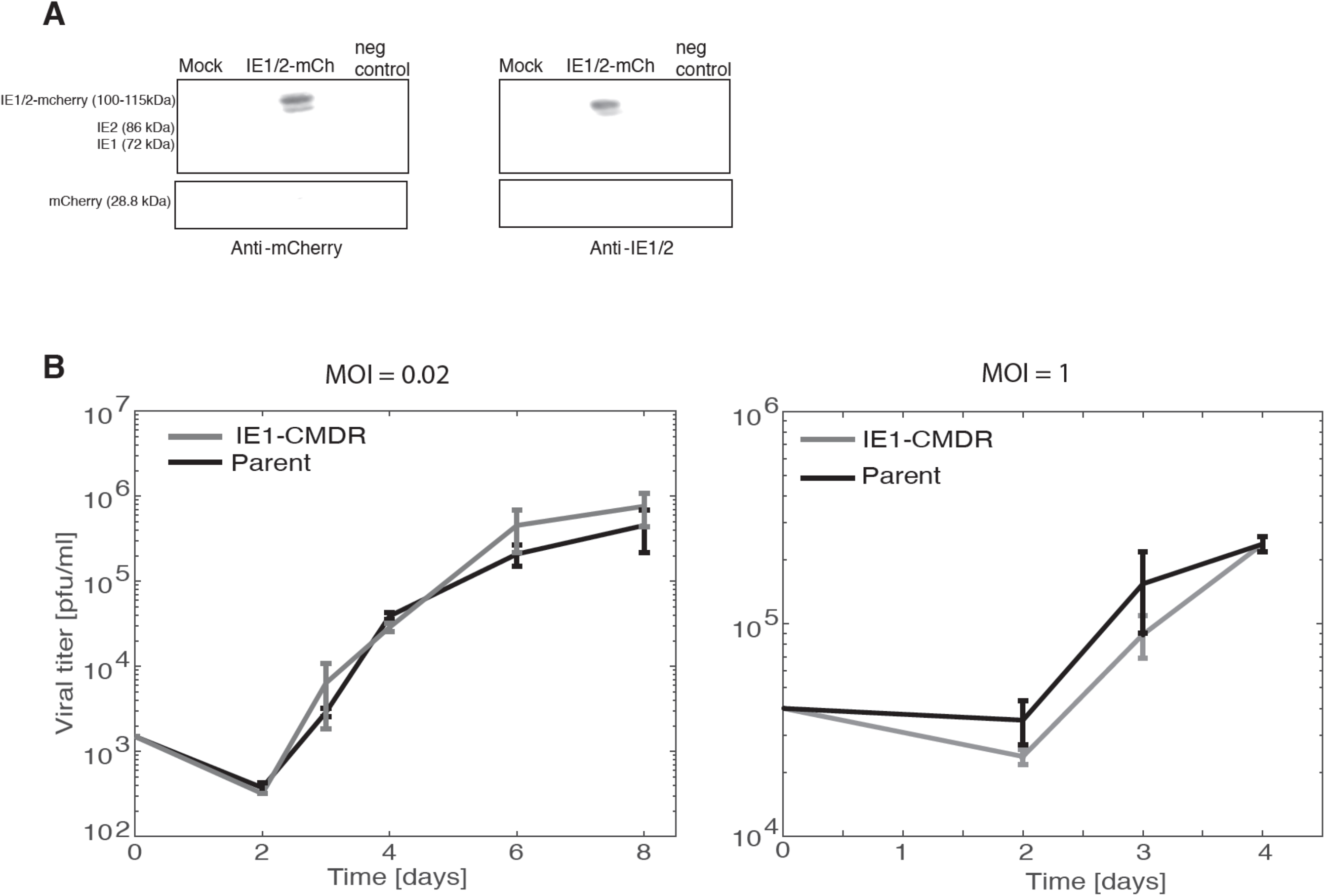
Fitness of IE1-CMDR virus is comparable to the parent virus. *(A)* Co-ip of IE1 and mCherry shows no free mCherry in IE1-CMDR infected ARPE-19 cells. Cells were either mock infected or infected with EYFP-pp71 virus at MOI = 1, immunoperccipitated with mCherry anti-body and stained with anti-mCherry (left panel) or anti-IE1/2 (right panel) antibodies. Immunoprecipitation of virus infected cell extract in the absence of mCherry antibody served as a negative control (right columns). See materials and methods for further details. (*B*) Viral titers from ARPE-19 cells infected with IE2-EYFP (black) or IE1-CMDR (grey) virus (average of two repeats, error bars denote standard deviation. Cells were infected at MOI=0.02 Left panel) or MOI=1 (right panel). Infections and titrations were done in medium containing 1 µM Shield-1.

**Figure S4:**
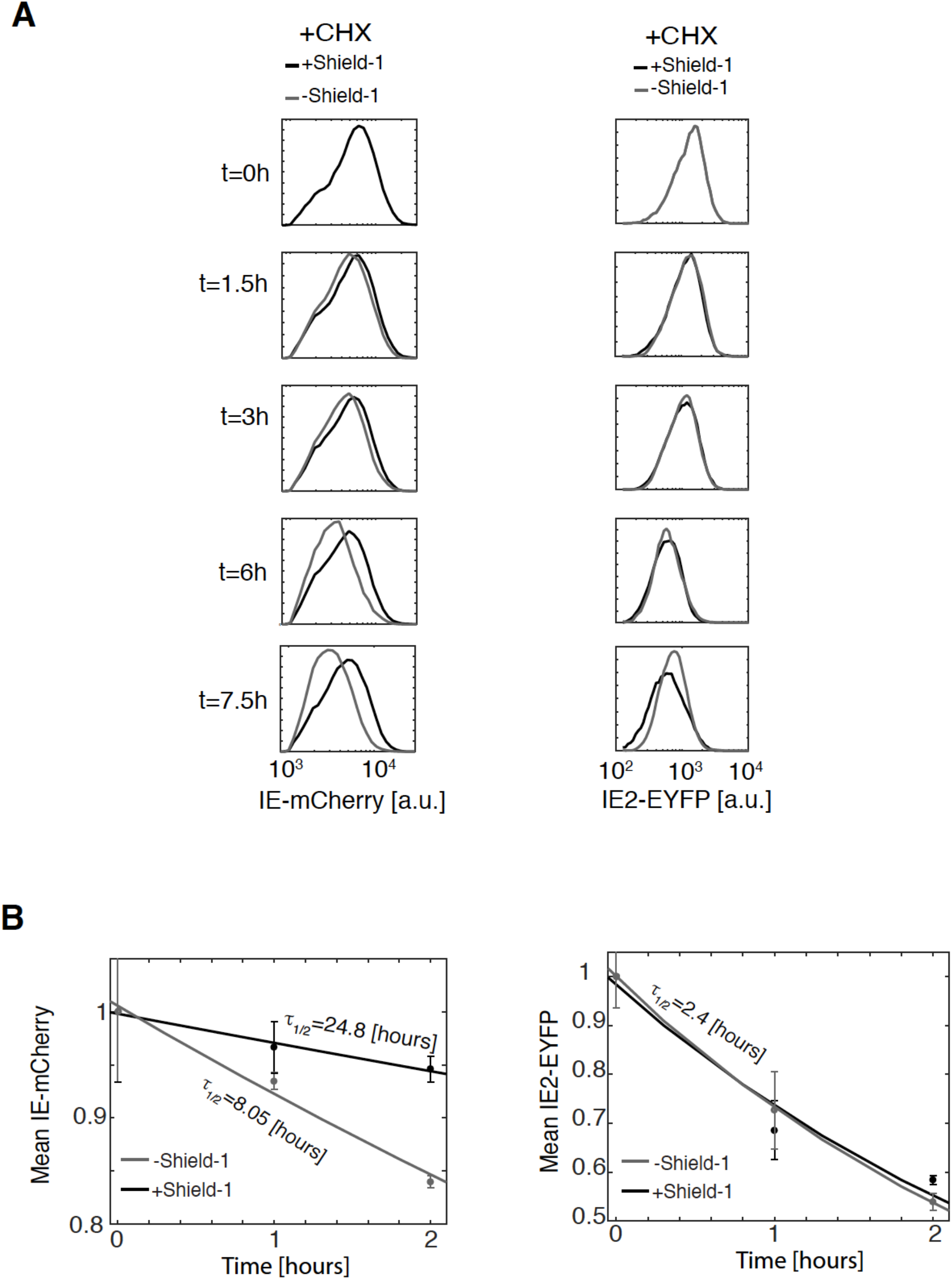
FKBP tag affects IE1 but not IE2 half-life. (*A*) 24 hours after infection with IE1-CMDR virus (MOI = 0.05), translation was blocked. IE-mCherry and IE2-EYFP levels were measured by flow-cytometry at 1 µM Shield-1 (dark grey) or without Shield-1 (light grey). (*B*) Quantification of IE-mCherry and IE2-EYFP decay reveals a three-fold difference in IE-mCherry rate and no difference in IE2-EYFP decay rate when CHX was added 15 hours post infection (average of two repeats, error bars denote standard deviation. Decay rate was calculated by fitting the data to an exponential decay model.

**Figure S5:**
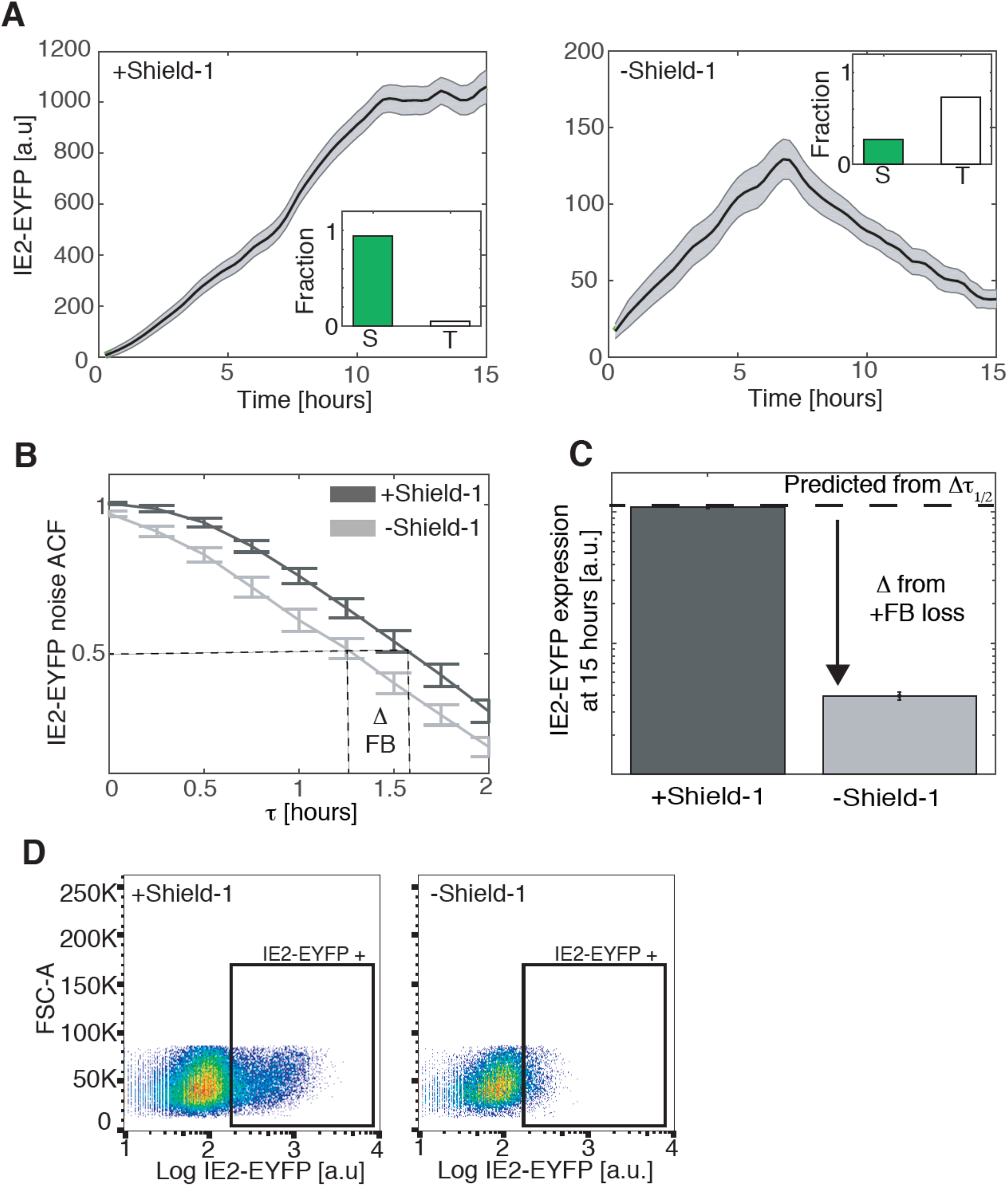
IE1 positive feedback sustains IE2 expression. (*A*) IE2-EYFP expression of cells infected with IE1-CMDR virus. Cells were cultured in medium with 1 µM Shield-1 (left panel, WT feedback, averaged over 119 cells) or without Shield-1 (right panel, attenuated feedback, averaged over 153 cells). Bold line denotes mean EYFP expression; grey shading denotes standard error. Cell trajectories were synchronized to the first detection of EYFP signal. In the inset is the fraction of cells with sustained (S) or transient (T) IE2-EYFP expression over three biological repeats. (*B*) Autocorrelation function (ACF) of IE2-EYFP in cells infected with IE1-CMDR virus in the presence of 1 µM Shield-1 (dark grey) or without Shield-1 (light grey). Shown is an average over a 100 cells; error bars denote standard error. (*C*) IE2-EYFP mean expression 15 hours after infection. Dashed line is the expected value from change in IE2 half-life; additional difference is due to the loss of IE1 positive feedback. (D) Flow cytometry of ARPE-19 infected with IE1-CMDR virus at MOI=0.2, 24 hours post infection. Cells were cultured in medium with 1 mM Shield-1 (left panel) or without Shield-1 (right panel). The change in mean fluorescence intensity of the population of IE2-EYFP positive cells (i.e., cells in the EYFP+ gate) is approximately 2-fold (shown in Figure 2K) whereas the change in IE2-YFP intensity for individual IE2-YFP+ cells (i.e., not gated) can be >2 fold (consistent with change shown above in panel A of this figure).

**Figure S6:**
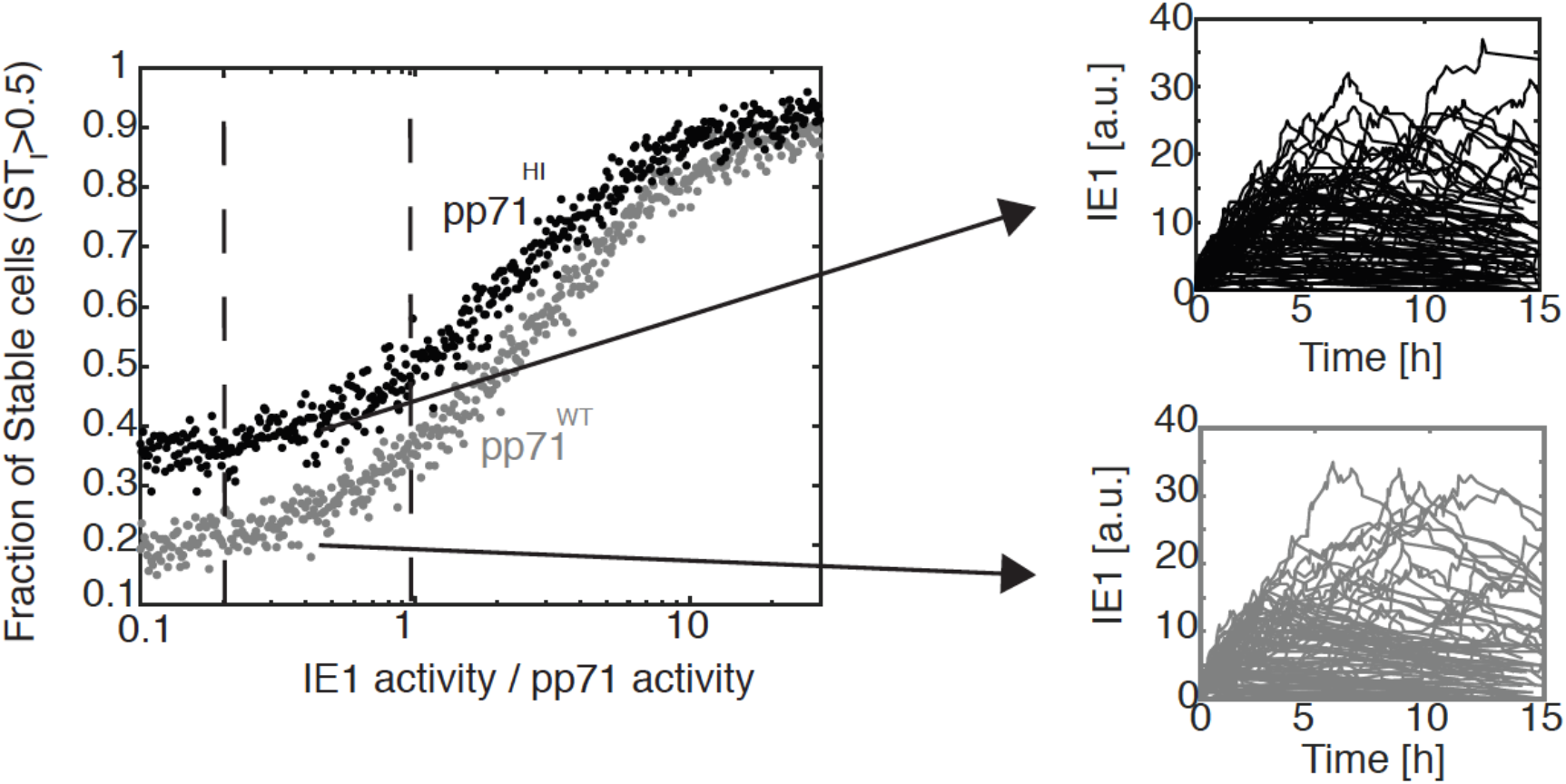
Stochastic simulations of the course grained pp71-IE1 model. A stochastic version of the course grained IE1-pp71 model was simulated using a Gillespie algorithm. The level of IE1 activity was varied over 2 orders of magnitude, decay rates of IE1 and pp71 were measured in Figures 1–2. The fraction of stable trajectories out of a 500 simulations is presented for each ratio of IE1/pp71 activity. For model parameters see Table S4.

### SUPPLEMENTARY TABLES

**Table S1:**
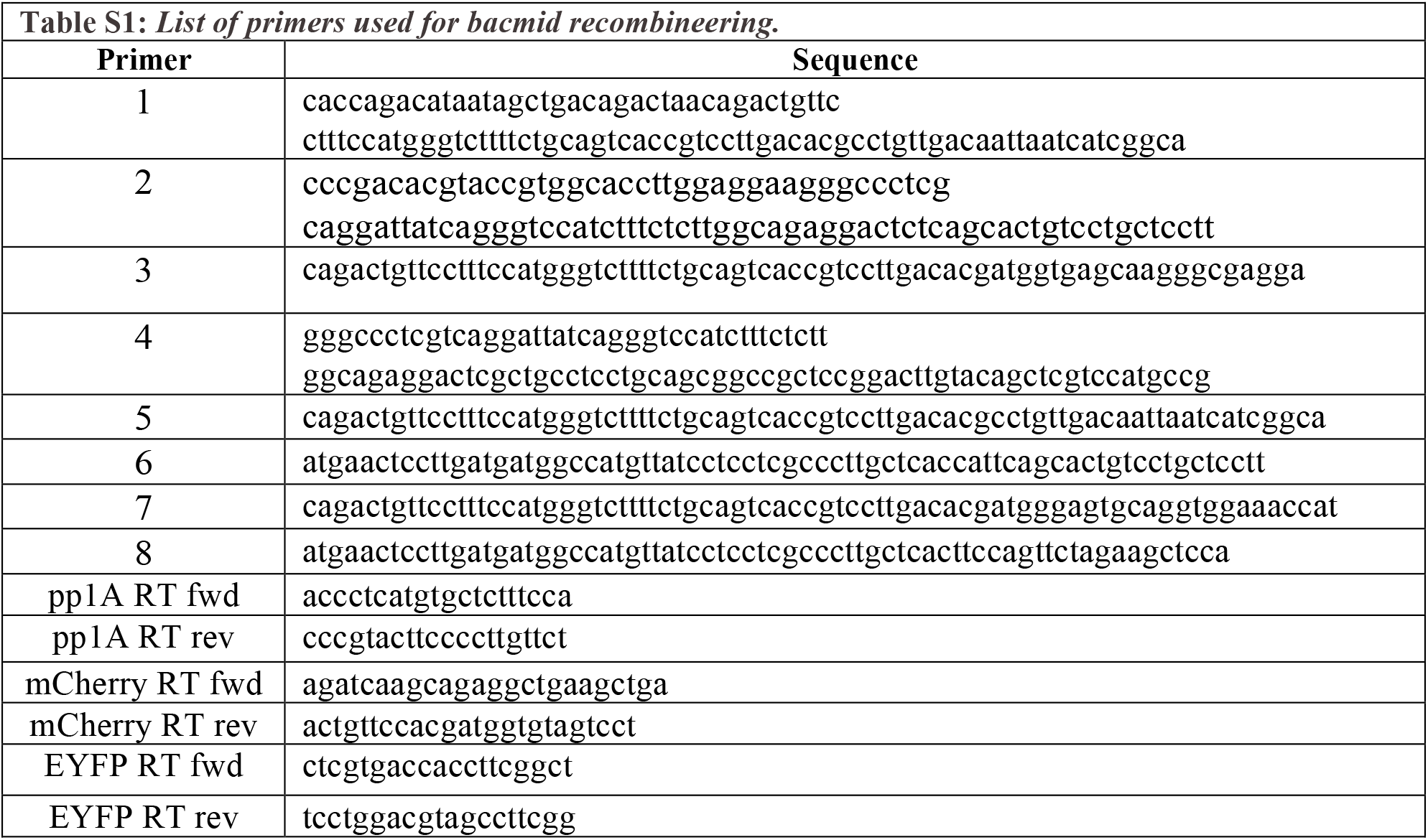
List of primers used for bacmid recombineering.

**Table S2:** Simulation parameters for Course grained model.

| Table S2: Simulation parameters for Course grained model. |  |
| --- | --- |
| Parameter | Value |
| $\alpha_0$ | 0.5 [nM/sec] |
| $\alpha_1$ | 82.4 [nM/hr] |
| $\alpha_2$ | 97.7 [nM/hr] |
| $\gamma_1$ | 0.03 - 0.1 [1/hr] |
| $\gamma_2$ | 0.11 [1/hr] |
| $h_1$ | 1.96 |
| $h_2$ | 2.03 |
| $k_1$ | 503 [nM] |
| $k_2$ | 478 [nM] |
| pp71 (t=0) | 200-1200 [nM] |
| IE1 (t=0) | 0 [nM] |

**Table S3:** Simulation parameters for detailed model (see Figure S2).

| **Table S3: Simulation parameters for detailed model (see Figure S2).** |  |
| --- | --- |
| **Parameter** | **Value** |
| $\alpha_0$ | 0.11 [nM/sec] |
| $\alpha_1$ | 13.2 [nM/hr] |
| $\alpha_2$ | 11 [nM/hr] |
| $\alpha_3$ | 0.1 [nM/hr] |
| $\alpha_4$ | 20 [nM/hr] |
| $\alpha_5$ | 19.4 [nM/hr] |
| $\gamma_1$ | 0.11 [1/hr] |
| $\gamma_2$ | 1.3 [1/hr] |
| $\gamma_3$ | 0.03 - 0.1 [1/hr] |
| $\gamma_4$ | 0.23 [1/hr] |
| $h_1$ | 1.22 |
| $h_2$ | 3.79 |
| $h_3$ | 6.4 |
| $k_1$ | 598 [nM] |
| $k_2$ | 402 [nM] |
| $K_3$ | 496 [nM] |
| $C$ | 0.7 |
| pp71 (t=0) | 200-1200 [nM] |
| pmRNA (t=0) | 0 [nM] |
| IE1 (t=0) | 0 [nM] |
| IE1 (t=0) | 0 [nM] |

**Table S4:** Simulation parameters for stochastic model (see Figure S6)

| **Table S4: Simulation parameters for stochastic model (see Figure S6)** |  |  |
| --- | --- | --- |
| **Reaction** | **Description** | **Parameter value** |
| $\text{MIEP}_{\text{OFF}} + \text{pp71} \rightarrow \text{MIEP}_{\text{ON}} + \text{pp71}$ | Activation of MIEP by pp71 | $K_1 = 0.01$ |
| $\text{MIEP}_{\text{OFF}} + \text{IE1} \rightarrow \text{MIEP}_{\text{ON}} + \text{IE1}$ | Activation of MIEP by IE1 | $K_2 = 0.001 - 0.1$ (varied) |
| $\text{IE1} \rightarrow 0$ | Decay of IE1 | 0.086 (from Figure 2) |
| $\text{pp71} \rightarrow 0$ | Decay of pp71 | 0.085 (from Figure 1) |
| $\text{MIEP}_{\text{on}} \rightarrow \text{MIEP}_{\text{on}} + \text{IE1}$ | Production of IE1 | 4.5 |
| $\text{MIEP}_{\text{on}} \rightarrow \text{MIEP}_{\text{off}}$ | Silencing of MIEP | 0.53 |

### SUPPLEMENTARY MOVIES

Movie S1: *Wild-type IE1 feedback*. Fluorescence time-lapse microscopy of IE-mCherry and IE2-EYFP in cells infected with IE1-CMDR virus (1µM Shield-1).

Movie S2: *Attenuated IE1 feedback*. Fluorescence time-lapse microscopy of IE-mCherry and IE2-EYFP in cells infected with IE1-CMDR virus (-Shield-1, attenuated IE1 feedback).

